# Temporal expectation modulates cortical dynamics of sensory memory

**DOI:** 10.1101/201756

**Authors:** Anna Wilsch, Molly J. Henry, Björn Herrmann, Christoph S. Herrmann, Jonas Obleser

**Affiliations:** Max Planck Research Group “Auditory Cognition”, Max Planck Institute for Human Cognitive and Brain Sciences, Leipzig, Germany; Experimental Psychology Lab, Center for Excellence “Hearing4all,” European Medical School, University of Oldenburg, Oldenburg, Germany; Department of Psychology, University of Lübeck, Lübeck, Germany

## Abstract

Increased memory load is often signified by enhanced neural oscillatory power in the alpha range (8–13 Hz), taken to reflect inhibition of task-irrelevant brain regions. The corresponding neural correlates of memory decay, however, are not yet well-understood. Here, we investigated auditory sensory memory decay using a delayed matching-to-sample task with pure-tone sequences. First, in a behavioral experiment we modeled memory behavior over six different delay-phase durations. Second, in a magnetoencephalography (MEG) experiment, we assessed alpha-power modulations over three different delay-phase durations. In both experiments, the temporal expectation for the to-be-remembered sound was manipulated, so that it was either temporally expected or not. In both studies, memory performance declined over time but this decline was less strong under a more precise temporal expectation. Similarly, patterns of alpha power in and alpha-tuned connectivity between sensory cortices changed parametrically with delay duration (i.e., decrease in occipito-parietal regions, increase in temporal regions). Notably, temporal expectation counteracted alpha-power decline in heteromodal brain areas (i.e., supramarginal gyrus), in line with its memory-decay counteracting effect on performance. Correspondingly, temporal expectation also boosted alpha connectivity within attention networks known to play an active role during memory maintenance. The present data outline how patterns of alpha power orchestrate sensory memory decay, and encourage a refined perspective on alpha power and its inhibitory role across brain space and time.

**Significance Statement:** Our sensory memories of the physical world fade quickly. We show here that this decay of sensory memory can be counteracted by so-called temporal expectation, that is, knowledge of when to expect the to-be-remembered sensory event (here, brief sound patterns). We also show that distinct patterns and modulations of neural oscillations in the “alpha” (8–13 Hz) range index both, the degree of memory decay, and any benefit from temporal expectation, both of which affect memory performance. Critically, spatially distributed cortical patterns of alpha power, with opposing effects in auditory vs. visual sensory cortices and alpha-tuned connectivity changes within supramodal attention networks, reflect the allocation of neural resources as sensory memory representations fade.

## Introduction

Working memory allows us to focus our attention on representations of perceptions that are no longer physically present (Baddeley, 2012). This ability is limited, though, by memory load and memory decay. Memory load reflects a capacity limit: The amount of information as well as a lack of precision of information demand memory capacity and must not exceed a certain limit in order to be stored (e.g., Luck and Vogel, 1997; van den Berg et al., 2012; Ma et al., 2014; Joseph et al., 2016). Memory decay refers to fading away of the memory representation over time (Brown, 1958; Posner and Keele, 1967). Neural oscillations in the alpha range (8–13 Hz), recorded using human electroencephalography (EEG) or magnetoencephalography (MEG), are modulated by manipulations of memory load. For example, alpha power increases when the number of items that a person is asked to hold in memory increases (Jensen et al., 2002; Busch and Herrmann, 2003; Leiberg et al., 2006; Obleser et al., 2012). However, it is less clear how neural oscillatory activity is related to memory decay. The current study examined the time course of alpha power as auditory information decayed from working memory.

Previous work on the neural correlates of memory decay suggests a reduction of neural responses during the “delay phase”, that is, the time during which information is held in memory before it can be reported or compared to another stimulus. Over the time of a memory-delay phase, single-cell activity in monkey prefrontal cortex decreases (Fuster, 1999), as does the BOLD response measured in posterior cortical regions in humans (Jha and McCarthy, 2000; for visual memory) and in temporal regions (Gaab et al., 2003; for auditory memory). Given the relationship between BOLD responses and cortical alpha power (Sadaghiani et al., 2010), we hypothesized that alpha power would also decrease over a memory delay phase.

One factor that has the potential to protect sensory information from decay during the delay phase is temporal expectation. Detection and discrimination are more accurate for temporally expected compared to unexpected stimuli (Coull and Nobre, 1998; Griffin et al., 2001; Nobre, 2001; Jaramillo and Zador, 2011), and temporally expected events contribute more strongly than unexpected events to perceptual evidence accumulation (Cravo et al., 2013). We have previously shown that temporal expectation reduces memory load for speech-in-noise, as indexed by improved memory performance for temporally expected stimuli (Wilsch et al., 2015a). Notably, this load reduction was accompanied by decreased alpha power during stimulus retention. Moreover, temporally expected distractors are more easily kept out of working memory than unexpected distractors, and this effect was also accompanied by increasing alpha power in anticipation of expected distractors (Bonnefond and Jensen, 2012). It is unclear, however, whether temporal expectation also has a beneficial effect on memory decay (see Kunert and Jongman, 2017).

Here we report the results of two experiments investigating the time course of decay of sensory memory (Cowan, 1984; Cowan et al., 1997; Nees, 2016). Auditory sensory memory enables integration of auditory information and preservation of information over brief periods of time (Schröger, 2007). We conducted a delayed pitch comparison procedure (e.g., Harris, 1952; Bachem, 1954; Bull and Cuddy, 1972; Keller et al., 1995) with two brief pure-tone sequences embedded in noise, separated by variable delay phases asking whether both sequences were same or different from each other. We made use of non-verbal stimuli to preclude rehearsal effects (Obleser and Eisner, 2009; Oberauer and Lewandowsky, 2013) and thus to keep any effects interpretable in terms of sensory memory.

Experiment 1 probed and modelled memory performance over six increasing delay phases. We addressed the question whether temporal expectation affects memory decay behaviorally. In order to assess how temporal expectation and memory decay interact at the neural level and specifically in terms of neural alpha (~8–13 Hz) oscillatory dynamics, Experiment 2 investigated their relationship using MEG. Alpha power modulations were assessed on the sensor level as well as by means of source analyses and functional connectivity.

## Methods

### Participants

Nineteen healthy right-handed participants (12 females; age range 23–33 years, median 25 years) took part in Experiment 1 (behavior and modelling). An independent sample of twenty healthy right-handed participants (10 females) ranging in age from 23 to 33 (median = 27) years took part in Experiment 2 (behavior and MEG recordings). All participants had self-reported normal hearing. Participants were fully debriefed about the nature and goals of the studies, and received financial compensation of 7 € per hour for their participation. The local ethics committee (University of Leipzig) approved of the studies, and written informed consent was obtained from all participants prior to testing.

### Experimental task and stimuli

The time course of an example trial is depicted in Figure 1A. On each trial, participants heard two pure-tone sequences (S1 and S2, see “Characteristics of sound stimuli”) and responded whether they were the same or different. These pure-tone sequences were embedded in noise, in order to increase perceptual load (Pichora-Fuller and Singh, 2006; van den Berg et al., 2012). Each trial began with the presentation of a fixation cross. After a brief pause (jittered between 0.75 s and 1.25 s), white noise and a visual cue were presented simultaneously. The visual cue indicated the onset time of the first sound (S1; see next paragraph) and remained on screen throughout the entire trial. Participants had to retain S1 in memory for a variable period of time. Then, a second sound (S2) was presented, and participants made a “same”/“different” judgment by pressing one of two buttons on a response box. The response was prompted approximately 1 second (jittered between 0.9 s and 1.1 m) after the presentation of S2. Finally, participants indicated their confidence in their “same”/“different” response on a 3-level confidence scale (“not at all confident”, “somewhat confident”, “very confident”). Trials were separated by an inter-trial interval of around 1 second (jittered: 0.75–1.25 s) that was free of stimulation or responses. See Figure 1A for an outline of a trial.

**Figure 1.**
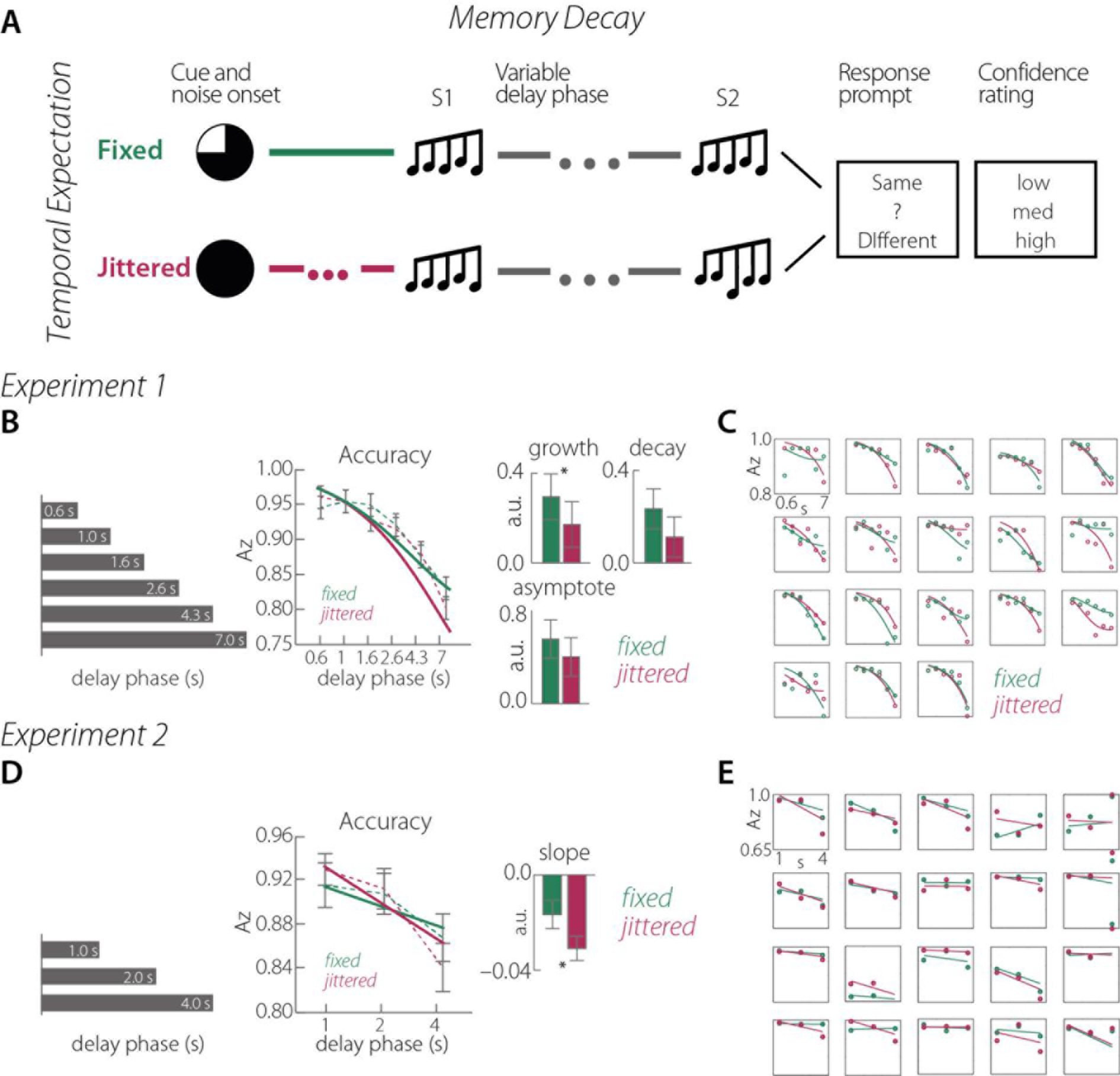
Experimental design and behavioral performance. **A.** Experimental design. The upper panel illustrates a “same” trial (S1 and S2 are the same) with a fixed onset time. The lower panel illustrates a “different” trial (S1 and S2 are different) with jittered onset time. The actual durations of the variable delay phases are specified in **B** and **D. B.** Memory performance in Experiment 1. The gray bars illustrate the six variable delay-phase durations from 0.6 s to 7.0 s (i.e., values in each bar). The line graph displays averaged memory performance in A_z_ (dotted lines) and the exponential fit (solid lines), both separately for fixed and jittered onset times; error bars indicate standard error of the mean of A_z_. The bar graphs show the average values for the estimated parameters “growth” and “decay”, as well as the asymptote, separately for fixed and jittered onset times. Error bars display the standard error of the mean. In all graphs, green refers to fixed and magenta to jittered onset times. The asterisk indicates the significant difference between fixed and jittered onset times. **C.** Single-participant exponential fits. Every single plot displays the exponential fit of one participant separately for fixed (green) and jittered (magenta) onset times. Dots display the actual performance data A_z_. **D.** Memory performance in Experiment 2. The gray bars illustrate the three variable delay phase durations from 1.0 s to 4.0 s (i.e., values in each bar). The line graph displays averaged memory performance in A_z_ (dotted lines) and the exponential fit (solid lines), both separately for fixed and jittered onset times; error bars indicate standard error of the mean of A_z_. The bar graph shows the average values for the estimated slope, separately for fixed and jittered onset times. Error bars display the standard error of the mean. In all graphs, green refers to fixed and magenta to jittered onset times. The asterisk indicates the significant difference between fixed and jittered onset times. **E.** Single-participant linear fits. Every single plot displays the linear fit of one participant separately for fixed (green) and jittered (magenta) onset times. Dots display the actual performance data A_z_.

#### Operationalization of memory decay and temporal expectation

Memory decay was manipulated by varying the time interval (delay phase) between S1 and S2. The aim of Experiment 1 was to fit an exponential decay function to memory performance across different delay-phase durations. That is why the delay-phase duration was varied logarithmically in six steps ranging between 0.6 and 7 s (i.e., 0.6, 1, 1.6, 2.6, 4.3, 7 s; see Figure 1B, left panel). In Experiment 2, delay phases were more coarsely sampled (1, 2, and 4 seconds; see Figure 1D, left panel).

Temporal expectation for S1 was manipulated by varying the S1-onset times relative to the presentation of a visual cue. Onset times were either fixed (i.e., S1 occurred 1.3 m after the onset of the visual cue) or jittered (i.e., S1 occurred after a duration drawn from a uniform distribution ranging between.9 s and 1.7 s, mean = 1.3 s.

#### Characteristics of the sound stimuli

All sound stimuli were sequences consisting of five pure tones; each pure tone had a duration of 40 ms resulting in a total sound duration of 200 ms (Watson et al., 1975). Sound stimuli were presented in standard-deviant pairs. For the standard stimulus, the middle (third) tone’s frequency was randomly selected on each trial from a uniform distribution ranging between 450 and 600 Hz. The second and fourth tones were independently assigned frequencies ±1–4 semitones (ST) with respect to the frequency of the middle tone, and the first and final tones were independently assigned frequencies ±4–7 ST with respect to the middle tone. Unique patterns were generated on each trial.

On half of the trials, a deviant stimulus was presented (i.e., “different” trials). For the deviant stimulus, the third and the fourth pure tone in the sequence were higher in frequency compared to S1. The third and fourth tones were both shifted up by the same amount (in ST; see Procedure). The exact standard-to-deviant-difference was adjusted for each participant individually (see “Procedure”). Each pure tone had an onset-and offset-ramp of 10 ms. On half of the trials, the standard stimulus was presented during the S1 interval, while the deviant stimulus was presented during the S1 interval the other half of the trials.

The noise masker was white noise. Sound sequences and noise were presented with a constant signal-to-noise ratio (SNR) of −17 dB. This SNR was determined via pilot testing to increase difficulty of the memory task but still allow all participants to perform the task.

### Procedure

Prior to the MEG measurement, participants were familiarized with the stimuli and task, and performed a few practice trials. Then, individual thresholds were estimated (i.e., the frequency difference between standard and deviant in the third and fourth pure tone position of the sound sequences). A custom adaptive-tracking procedure was utilized that yielded a frequency difference corresponding to memory performance falling between 65% and 85% correct responses.

In Experiment 1, participants completed 360 trials in 10 blocks of 36 trials each. In Experiment 2, brain activity was recorded with MEG during the performance of 396 trials completed in 12 blocks of 33 trials each. The manipulation of S1-onset time (fixed, jittered) was kept constant within a block, and participants were informed at the start of each block about the type of temporal cue they would receive on each trial. Delay-phase durations (0.6–7 seconds, and 1-, 2-, 4-seconds, for Experiment 1 and 2 respectively) were equally distributed within blocks. The order of trials within a block and order of blocks were randomized for each participant. Button assignments were counterbalanced across participants, such that half of the participants indicated that the first and the second sound were the same using the left button, and half did so with the right button.

The testing took approximately 2.5 hours per participant and was conducted within one session. The overall session including practice blocks and preparation of the MEG setup took about 4 hours.

### Modelling of behavioral data in Experiment 1

#### Data analysis

The crucial measure for memory decay was the performance measure A_z_, a non-parametric performance measure derived from confidence ratings. Confidence ratings were used to construct receiver operating characteristic (ROC) curves (Macmillan and Creelman, 2004) for each condition, and ROC curves were used to derive A_z_. A_z_ can be interpreted similarly to proportion correct. A_z_ was computed for each of the twelve conditions (temporal expectation, 2, × memory decay, 6), allowing us to estimate memory decay as a function of delay-phase duration separately for fixed and jittered onset times. One participant had to be excluded from this analysis because the participant did not make use of the entire confidence rating scale in at least two experimental conditions; A_z_ could not be computed for these data points. Another participant presented the same behavior but only in one condition. Here, the missing A_z_ value was interpolated by calculating the mean of the two adjacent conditions.

We fitted Equation 1 (Glass and Mackey, 1988) to A_z_ scores as a function of delay-phase duration:

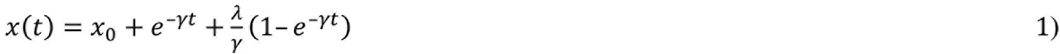

where *t* is equal to time (i.e., delay-phase duration) and *x*_0_ corresponds to the intercept. This specific function contained a term describing decay, *ϒ*, and an additional term describing growth, *λ*. This function has the advantage (as compared to simple decay functions; e.g., Wickelgren, 1969; Rubin and Wenzel, 1996) that it takes the nature of physiological systems into account. That is, it assumes that in physiological systems activation declines as new activation simultaneously arises: during working memory retention, the memory representation decays over time, but allocation of cognitive resources can counteract that decay. Note that 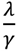 indicates the function's asymptote.

The initial parameters for the function fits were as follows: *x*_0_ = 0, *γ* = 0, and *λ* = 0, where *x*_0_ was bound between zero and one, and *γ* and *λ* were bound between zero and infinity. The model fit was computed with the lsqcurvefit function with Matlab (version 8.2, Optimization Toolbox) that allowed for 1000 iterations.

In addition, we also fitted a decay-term-only model (i.e., first term: x(t) = (x_0_ + e^−γt^)). The decay-only model is more parsimonious and more commonly used to estimate memory decay (Peterson and Peterson, 1959; Wickelgren, 1969). To determine which one of these two models represented the memory performance data best, we calculated the Bayesian information criterion ( BIC; Schwarz, 1978) for both model fits, as well as for fixed and jittered onset times separately. Note that the BIC penalizes for more parameters and allows for an equitable comparison of goodness-of-fit of both models (smaller is better). We averaged the BICs across fixed and jittered onset times separately for each function. 17 out of 18 participants had a lower BIC for the full model (Equation 1) than the decay-only model (t(17) = 4.75, p < 0.001) indicating an overall better fit by the former model. Therefore, all further analyses were conducted on the parameters resulting from the fit of the complete Equation 1. Four of the participants were excluded from the t-tests, because R2, an indicator for goodness of the model fit, of their fitted models was smaller than 0.3 (see Figure 1C for individual model fits). The average R2 values for the fixed and jittered conditions, respectively, were 0.66 (sd = 0.31; 0.80, sd = 0.13 without excluded participants) and 0.72 (sd = 0.25; 0.81, sd = 0.16 without excluded participants).

After the fitting of the function, the resulting parameters *x*_0_, *γ*, and *λ* for jittered and fixed onset times as dependent variables were assessed with a multivariate ANOVA. This allowed us to test whether there is a global difference between jittered and fixed onset times. Subsequently, the parameters *γ*, *λ*, and x_0_ were tested for differences between fixed and jittered onset times with univariate repeated-measures ANOVAs, in order to determine whether memory decay was less strong when S1-onset times were predictable.

### Data recording and analysis in Experiment 2

Participants were seated in an electromagnetically shielded room (Vacuumschmelze, Hanau, Germany). Magnetic fields were recorded using a 306-sensor Neuromag Vectorview MEG (Elekta, Helsinki, Finland) with 204 orthogonal planar gradiometers and 102 magnetometers at 102 locations. Two electrode pairs recorded a bipolar electrooculogram (EOG) for horizontal and vertical eye movements. The participants' head positions were monitored during the measurement by five head position indicator (HPI) coils. Signals were sampled at a rate of 1000 Hz with a bandwidth ranging from direct current (DC) to 330 Hz.

The signal space separation method was applied offline to suppress external interferences in the data, interpolate bad channels, and to transform individual data to a default head position that allows statistical analyses across participants in sensor space (Taulu et al., 2004).

Subsequent data analyses were carried out with Matlab (The MathWorks Inc., Massachusetts, USA) and the FieldTrip toolbox (Oostenveld et al., 2011) using only trials to which correct responses were provided (“correct trials”). Analyses were conducted using only the 204 gradiometer sensors, as they are most sensitive to magnetic fields originating directly underneath the sensor (Hämäläinen et al., 1993). The continuous data were filtered offline with a 0.5-Hz high pass filter, specifically designed to provide a strong suppression of DC signals in the data (>140 dB at DC, 3493 points, Hamming window; e.g., Ruhnau et al., 2012).

Subsequently, trial epochs ranging from −1.5 to 11.5 s time-locked to the onset of S1 were defined. The use of long epochs prevented windowing artifacts in the time-frequency analysis; the intervals analyzed statistically were shorter (see below). Epochs were low-pass filtered at 80 Hz and subsequently down-sampled to 200 Hz.

Epochs with strong artifacts were rejected when the signal range at any gradiometer exceeded 800 pT/m. Independent component analysis (ICA) was applied to the epochs in order to reduce artifacts due to eye blinks and heartbeat. Following ICA, remaining epochs were rejected when the signal range within one epoch exceeded 200 pT/m (gradiometer) or 100 μV (EOG). Additionally, trials were rejected manually for which variance across sensors was deemed high relative to all others (per participant, per condition) based on visual inspection. For further analysis, each trial was time-locked at two different points, i.e., all trials were time-locked to the first stimulus (t = 0 s at S1 onset) and to the second stimulus (t = 0 s at S2 onset) for separate analyses. This was because different trials had different delay phase durations so that trials time-locked to S1 were not always time-locked to S2.

### Spectral analysis

The focus of the spectral analyses was on the set of trials time-locked to S2, allowing for analyses related to the end of the delay phase. For each trial, a 0.7-s segment was extracted (−.8 to −0.1 s time-locked to S2 excluding evoked responses due to S1 sound presentation), multiplied with a Hann taper, and the power between 8–13 Hz was computed using a fast Fourier transform (FFT).

For illustration purposes only, we also computed time-frequency representations (TFRs) of trials that were time-locked to S1. Time-frequency analysis was conducted on trial epochs ranging from −2.0 to 7.6 s for each trial (with 20-ms time resolution) for frequencies ranging between 0.5 Hz to 20 Hz (logarithmically spaced, in 20 bins). Single-trial time-domain data were convolved with a Hann taper, with an adaptive width of two to four cycles per frequency (i.e., 2 cycles for 0.5−1.6 Hz, 3 cycles for 1.9−9.2 Hz, and 4 cycles for 11.1-20 Hz). The output of the analysis was complex Fourier data. For further analyses, power (squared magnitude of the complex-valued TFR estimates) was averaged across single trials. Inter-trial phase coherence (ITPC) was computed based on the complex Fourier data (Lachaux et al., 1999). ITPC is the magnitude of the amplitude-normalized complex values averaged across trials for each time-frequency bin per channel and experimental condition (Thorne et al., 2011).

Next, FFT power spectra as well as TFRs were averaged across gradiometers in each pair. This procedure resulted in one value for each time point (TFRs only), frequency bin and sensor position of every single trial for each delay-phase condition and onset-time condition.

### Source localization

In order to estimate the origin of sensor-level alpha-power, source localizations were computed based on individual T1-weighted MRI images (3T Magnetom Trio, Siemens AG, Germany). Topographical representations of the cortical surface of each hemisphere were constructed with Freesurfer (http://surfer.nmr.mgh.harvard.edu) and the MR coordinate system was co-registered with the MEG coordinate system using the head-position indicators (HPIs) and about 100 additional digitized points on the head surface (Polhemus FASTRAK 3D digitizer). For forward and inverse calculations, boundary element models were computed for each participant using the inner skull surface as volume conductor (using the MNE toolbox; https://martinos.org/mne/). Individual mid-gray matter surfaces were used as source model by reducing the approximately 150,000 vertices needed to describe single hemispheres to 10,242 vertices.

The beamformer approach (DICS, dynamic imaging of coherent sources; Gross et al., 2001) was used to project alpha power (−0.8 to −0.1 s time-locked to S2-onset) to source space. To this end, a multitaper FFT centered at 11 Hz (±2 Hz smoothing with three Slepian tapers; Percival and Walden, 1993) was computed. A complex filter was calculated based on the data of all delay-phase and onset-time conditions (Gross et al., 2001; Schoffelen et al., 2008). Single-trial complex FFT data were then projected through the filter, separately for each condition providing a power value for each frequency bin in the alpha range at each vertex.

Neural activity was spatially smoothed across the surface (vertices) using an approximation to a 6 mm FWHM Gaussian kernel (Han et al., 2006). Individual cortical representations were transformed to a common coordinate system (Fischl et al., 1999b), and finally morphed to the pial cortical surface of the brain of one participant for display purposes (Fischl et al., 1999a).

#### Functional connectivity analyses

In order to attain a better understanding of the functional role of alpha power in memory decay, specifically for alpha power emerging from left superior temporal gyrus (STG, MNI [−50, −17, −8]; see below), connectivity analyses between cortical sources were computed. A whole-brain approach was adopted to find brain areas that were functionally connected with left STG based on the basis of the phase-locking value (PLV; Lachaux et al., 1999; see also Keil et al., 2014). Fourier spectra from 8–13 Hz were calculated in the time window time-locked to S2 (−0.8 to −0.1 s) and multiplied by the previously calculated common DICS filter (see above). Then, single-trial complex Fourier spectra were transformed into angle values and the circular distance between each vertex and STG was calculated for each trial. Finally, the PLV, i.e., the resultant vector length of the circular distance, was calculated across trials at each vertex. The greater the PLV at a vertex the greater the phase coherence between this vertex and left STG.

### Statistical analysis

#### Memory performance

Analogous to Experiment 1, memory performance for each condition was indexed by A_z_ (see Figure 1D). Since in Experiment 2 only three different delay-phase durations were employed instead of six, we were only able to compute a linear fit across these durations. Hence, memory decay was estimated by regressing A_z_ on the delay phase durations of 1-, 2-, and 4 seconds. The impact of temporal expectation on memory decay was measured by comparing the slopes of the linear fit for fixed and jittered S1-onset times using a paired-samples t-test. Response times are not reported because responses were cued and thus do not provide valid information about costs and benefits of the experimental manipulations.

#### Sensor level analyses

Statistical analyses were only conducted on the FFT power spectra (−0.8 to −0.1 time-locked to S2). Analyses were conducted according to a multi-level approach. On the first (single-subject) level, we regressed alpha power on the delay phase durations (1, 2, 4-s) similar to the regression of memory performance (A_z_) on delay phase duration (see above). To test the parametric modulation of memory decay, the FieldTrip-implemented independent-samples regression t-test was performed (Maris and Oostenveld, 2007). The regression t-test provides the regression b-coefficient (i.e., slope of the modulation) for each frequency bin at each of the 102 sensor positions indicating the strength of the tested contrast. Here, in order to test for a linear relationship between alpha power and delay phase duration, contrast coefficients were selected corresponding to the actual delay-phase duration in seconds (i.e., 1, 2, 4). To test whether temporal expectation had an impact on this relationship, the same contrast was calculated for fixed and jittered onset times separately.

For the statistical analyses on the second (group) level, b-values resulting from the first-level statistics testing the parametric modulations of alpha power by the delay phase were tested against zero. In addition, to test whether the delay-phase modulation in the fixed condition differs significantly from the modulation in the jittered condition, b-values attained for each of the onset-time conditions separately were tested against each other. The tests against zero as well as the tests contrasting fixed and jittered conditions were conducted with FieldTrip’s dependent sample t-test using cluster-based permutation tests. The cluster test corrects for multiple comparisons resulting from testing each frequency-sensor combination. All cluster tests were two-tailed and were thus considered significant when p < 0.025.

We also tested for correlations between alpha power and memory performance (A_z_), averaging over experimental conditions, with a multi-level cluster test. On the first level, each participant's six A_z_ values (2 temporal-expectancy conditions × 3 delay phases) were correlated with the corresponding alpha-power values. On the second level, first-level correlation values were fisher's z transformed and tested against zero with a dependent samples cluster-based permutation t-test.

#### Source level analyses

Statistical analyses for source-projected alpha power as well as for PLVs reflecting functional connectivity between left STG and any other vertex were conducted with the same approach. The aim was to test whether either variable (alpha power or PLV) was modulated by delay-phase duration and whether this modulation was affected by temporal expectation.

Contrasts were calculated for each vertex separately. In order to test for a linear relationship of memory decay and alpha power in source space, source projected alpha power and the delay-phase duration (1, 2, 4 s) were z-transformed on a single-subject level. Then the delay phase duration served as a regressor and was fitted to the source power to test for a linear relationship of alpha power and delay-phase duration. The same approach was applied to test for effects of functional connectivity: PLVs were z-transformed and z-transformed delay-phase duration values were fitted to these PLVs.

The resulting regression coefficients at each individual vertex from both contrasts were then morphed onto a common surface in MNI space, respectively (Freesurfer average brain; Fischl et al., 1999b). For the interaction of temporal expectation and memory decay, the same linear regression was applied to the same data again but separately for each temporal-expectation condition. On the group level, regression coefficients of each contrast were tested against zero or fixed-onset-time coefficients were tested against jittered-onset-time coefficients, respectively, with vertex-wise t-tests. The resulting t-values were z-transformed and displayed on the average brain surface with contrast dependent uncorrected vertex-wise threshold of |z| ≥ 1.96 (Sohoglu et al., 2012).

Then, brain regions that showed statistical effects were identified by extracting the MNI-coordinate of the greatest z-value within one area of interest. Areas of interest were identified by visual inspection. The MNI coordinate was then used to identify the specific brain region using the MNI structural atlas.

#### Correlation of alpha power and A_z_ in source space

Analogous to the analyses on the sensor level, the correlation of source projected alpha power and A_z_ was calculated by correlating A_z_ with alpha power within condition at each vertex point. Here as well, the Fisher's z-transformed correlation values were tested against zero with vertex-wise t-tests. The resulting t-values were z-transformed and displayed on the average brain surface with an uncorrected vertex-wise threshold of |z| ≥ 1.96.

## Results

In the present study, we investigated whether and how temporal expectation ameliorates the decay of sound representations in sensory memory. Participants were asked to retain a sound in memory for a delay phase that varied in duration from trial to trial and to judge whether that sound was the same or different from a sound presented following the delay phase. We focused on behavioral performance as well as on neural oscillatory activity in the alpha frequency band.

### Experiment 1: Behavioral modelling of memory decay

In Experiment I, we estimated a “forgetting curve” based on fits of an exponential-decay function to A_z_ values as a function of delay-phase duration. Fits were conducted separately for fixed and jittered S1-onset times in order to assess the effect of temporal expectation on memory decay. In line with the broad literature on sensory memory decay, A_z_ declined with longer delay-phase durations. Interestingly, performance decayed differently for jittered and fixed onset times (Figure 1B). The two functions (jittered and fixed) show that for delay-phases up to one second, memory performance was the same following fixed and jittered onset times, whereas for longer delay phases, performance declined less severely following fixed compared to jittered onset times. Figure 1C displays the single-subject fits of the decay function.

A multivariate ANOVA showed that the estimated parameters decay factor, growth factor, and intercept (Wilk's approximated F(3,11) = 3.81, p = 0.043) differed for fixed versus jittered S1-onset times. Subsequent univariate tests on all parameters separately revealed that there was a trend-level effect of the decay factor *γ* (F(1,13) = 3.68, p = 0.077; Figure 1B). The univariate test on the growth factor, *λ*, showed that growth over delay-phase duration was significantly greater for fixed than for jittered onset times (F(1,13) = 4.95, p = 0.044; Figure 1B), converging with the test on the decay factor (y) that A_z_ declines faster after jittered than after fixed onset times. The univariate test on the intercept x**0** did not show a difference between onset times (F(1,13) = 0.04, p = 0.84). Next, we tested both asymptotes separately against 0.5, corresponding to memory performance at chance level. The asymptote parameter estimate corresponding to fixed onset times was significantly larger than chance (i.e., 0.5) as shown by a 95−% confidence interval (CI) of [0.52; 0.82], whereas the asymptote after jittered onset times did not differ from 0.5 [95% CI 0.26; 0.64] (Figure 1B). However, fixed and jittered asymptotes did not differ significantly from each other (t(13) = 1.47, p = 0.164). Thus, while memory performance declines to chance level following jittered onset times for longer delays, fixed onset times counteract this decline in memory performance.

### Experiment 2: Linear effects of memory decay and temporal expectation on behavioral performance

In line with the findings of Experiment 1, the comparison of the single-subject slopes of fixed and jittered onset times revealed that sensitivity of sensory memory performance (as indicated A_z_) after jittered onset times decayed faster than after fixed onset times (t(19) = 2.72, p = 0.013, see Figure 1D; see Figure 1E for single-subject linear fits).

### Experiment 2: Effects of memory decay and temporal expectation on alpha power

We were interested in how memory decay was affected by temporal expectation, and how this relationship was related to alpha-power modulation. Figure 2A (upper panel) illustrates overall power for all frequency bands (5–20 Hz) time-locked to the onset of S1 (averaged across all channels). Figure 2B presents the time-course of alpha power averaged across trials for each condition separately. Following S1, alpha power increases until the earliest occurrence of S2 (i.e., shortest delay phase of 1 second) and then decreases slowly. Inter-trial phase coherence (ITPC; Figure 2A, lower panel) is increased time-locked to the visual cue and the auditory events. Apart from the cue-related response, the ITPC peak frequency is below the alpha range for sound-related responses. In what follows, we will focus on alpha power.

**Figure 2.**
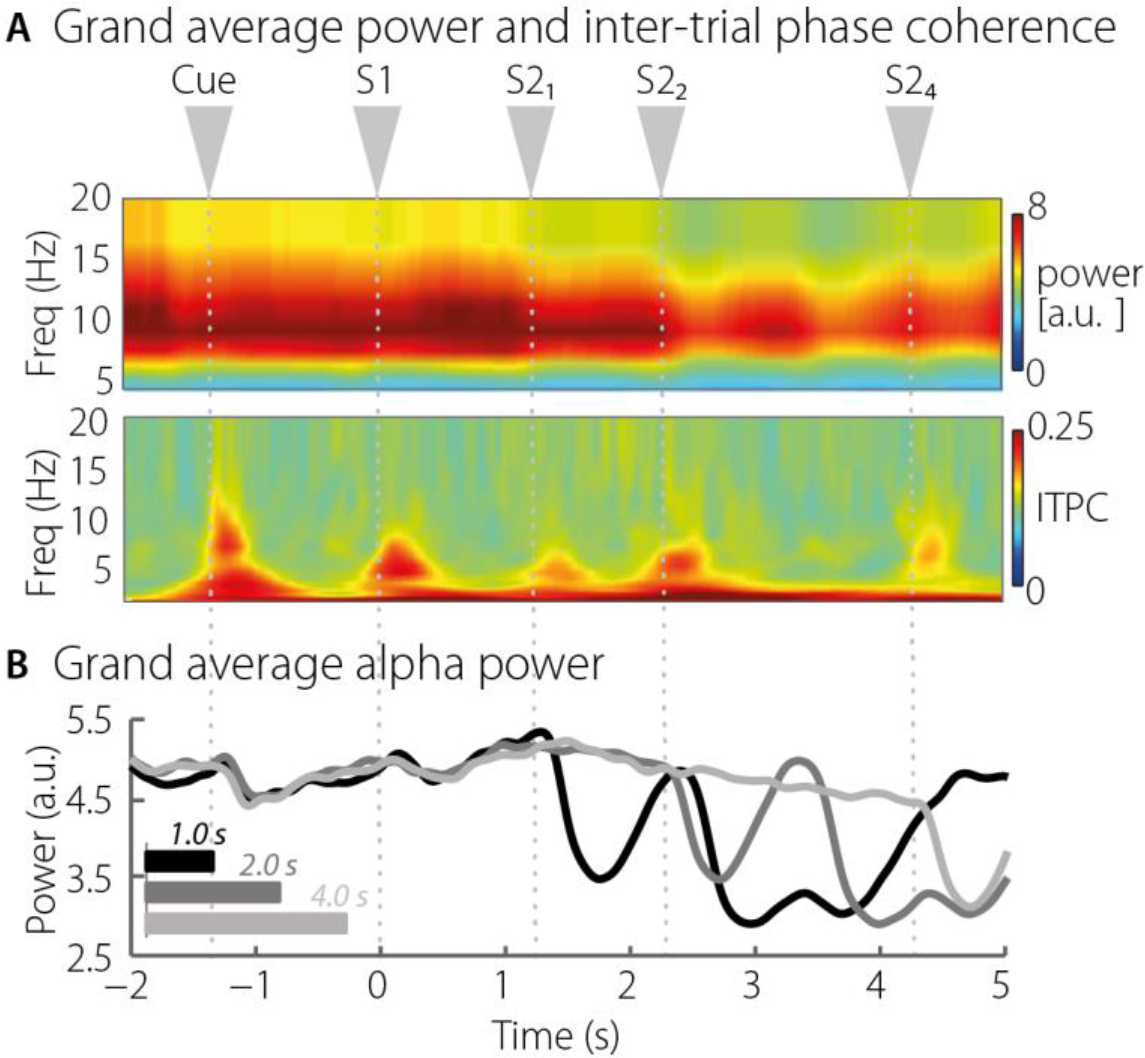
Time-Frequency grand averages power and phase coherence. **A.** Upper panel: Grand-average power 5-20 Hz averaged across all sensors. Gray arrows on top indicate stimulus occurrence times. S1 refers to the to-be-remembered stimulus. S2 refers to the second stimulus. The index indicates the corresponding delay-phase duration in seconds. Lower panel: Grand average of inter-trial phase coherence 5-20 Hz averaged across all sensors. **B.** Alpha power (8-13 Hz) grand-average across channels per delay-phase duration.

We investigated alpha-power changes as a function of delay phase (−0.8 to −0.1 s time-locked to S2, compare Figure 1) and whether the relationship between delay-phase duration and alpha power was modulated by temporal expectation.

The first-level b-coefficients resulting from the linear regression of alpha power on delay-phase duration were tested against zero on the group level. B-coefficients were significant<u>ly</u> smaller than zero in a broad posterior, negative cluster (p < 0.0001; see Figure 3A, upper panel) indicating that alpha-power decreased with longer delay-phase duration. A second cluster test contrasting the b-coefficients of the fixed onset-time condition with the b-coefficients of the jittered onset-time condition showed that temporal expectation also had an impact on alpha power: alpha power decreased less with increasing delay-phase durations following fixed onset times compared to jittered onset times (left-posterior positive cluster, p = 0.025; see Figure 3B, upper panel).

#### Source localization of alpha power modulations

Source localization was computed to identify the brain regions underlying the reported alpha-power effects on the sensor level. The effect of delay-phase duration on alpha power was localized at posterior and temporal sites. The negative peak indicating a decrease of alpha power with increasing delay-phase duration emerged from left primary visual cortex (V1, [MNI: −5, −88, 11]). In addition to the negative cluster on the sensor level, source localization reveals a positive linear relationship between alpha power and delay-phase duration emerging from left STG ([MNI: −50, −17, −8]). Z-transformed effects in source space and z-values greater than 1.96 for each delay-phase condition averaged across vertices around the peak effect in left V1 and left STG are illustrated in line graphs of Figure 3A (lower panel).

The differential effect of temporal expectancy on alpha power during the delay phase originated most prominently from the left supramarginal gyrus (SMG, [MNI: −54, −37, 32]) and right V1 ([MNI: 14, −80, 13]). Z-values greater than 1.96 are illustrated in Figure 3B, lower panel.

**Figure 3.**
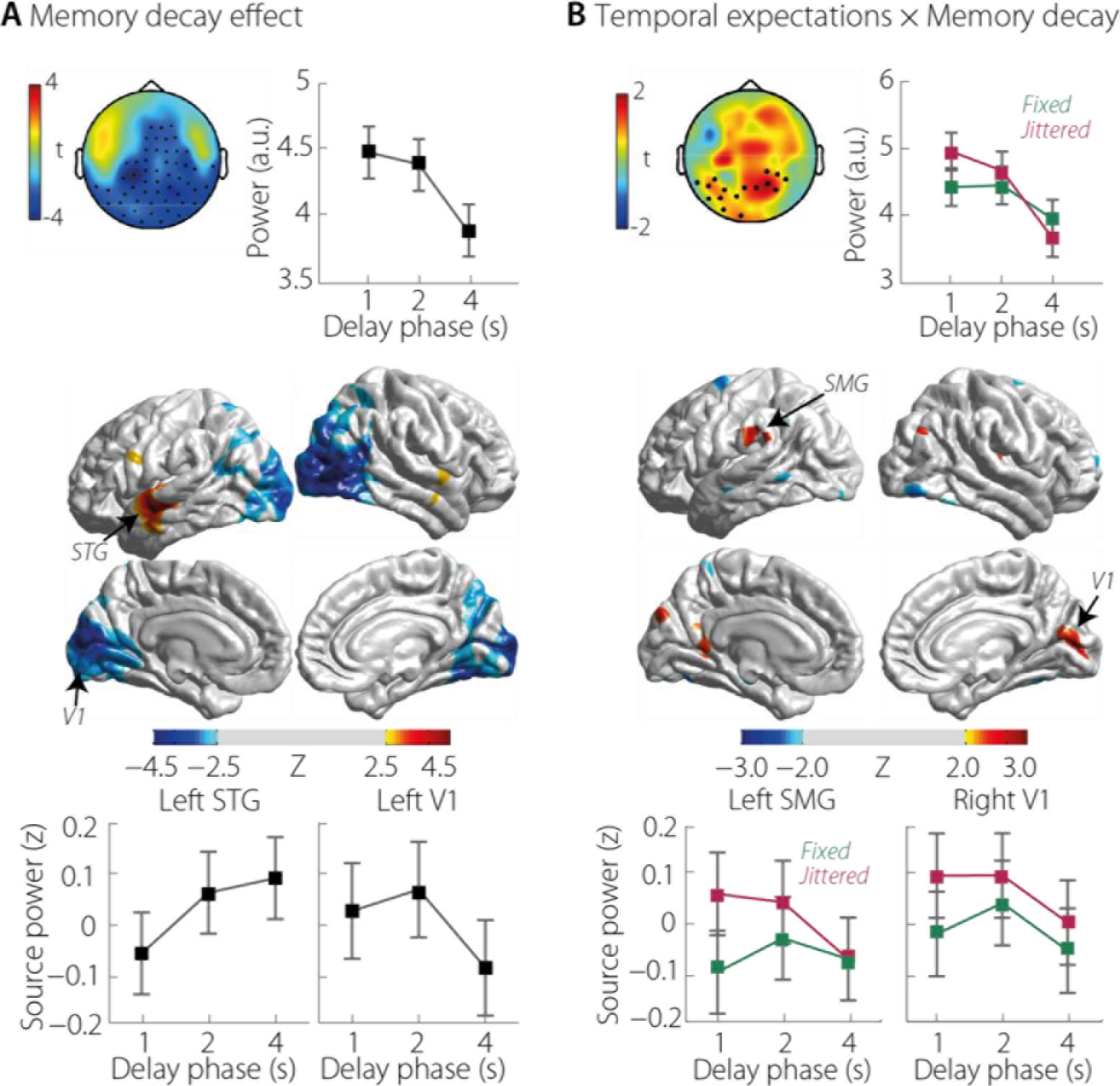
Condition effects in alpha power. **A.** Effect of memory decay (1, 2, 4 s delay phase). Upper panel: Topographies of the t-values of the linear fit of alpha power on delay-phase duration on the sensor level. Marked channels present the significant cluster. The line graph represents alpha power extracted from the displayed channels. Lower panel: Source projected linear fit of alpha power on delay-phase duration. Z-transformed t-values are displayed with a threshold of |z| ≥ 1.96. Line graphs display delay-phase activity drawn from and averaged across the vertices presenting peak activity around left STG and left V1. All error bars show within-subject standard error. **B.** Impact of temporal expectation on memory decay. Upper panel: Topographies of the t-values of the impact of onset-time condition on the linear fit of alpha power on delay-phase duration on the sensor level. Marked channels present significant cluster. Line graphs represent alpha power extracted from the displayed channels. Lower panel: Source projected difference between fixed and jittered onset times of the linear fit of alpha power on delay-phase duration. Z-transformed t-values are displayed with a threshold of |z| ≥ 1.96. Positive z-values indicate that jittered onset times have a steeper slope than fixed onset times. Line graphs display condition-wise activity drawn from and averaged across the vertices presenting peak activity around left SMG and right V1. All error bars display within-subject standard error.

#### Alpha power predicts behavioral performance

In a final analysis, we aimed to relate the observed modulation of memory performance (i.e. A_z_) to the alpha-power modulations. We correlated A_z_ and alpha power across all conditions by means of a cluster test, which revealed a centrally distributed positive cluster (p = 0.006; Figure 5A). Figure 5B illustrates the source projections of the correlation effect and figure 5C displays the single-subject correlations between A_z_ and source alpha drawn from left ACC. During the delay phase, the positive correlation of alpha power and A_z_ emerged from left anterior cingulate ([MNI: −2, 2, 38]), bilateral postcentral gyrus ([MNI: 28, −34, 70; MNI: −4, −9, 56]), and bilateral occipital cortices ([MNI: 7, −64, 62; MNI: −7, −86, 2]). A negative correlation between alpha power and A_z_ emerged from left STG ([MNI: −55, −10, −37]).

**Figure 4.**
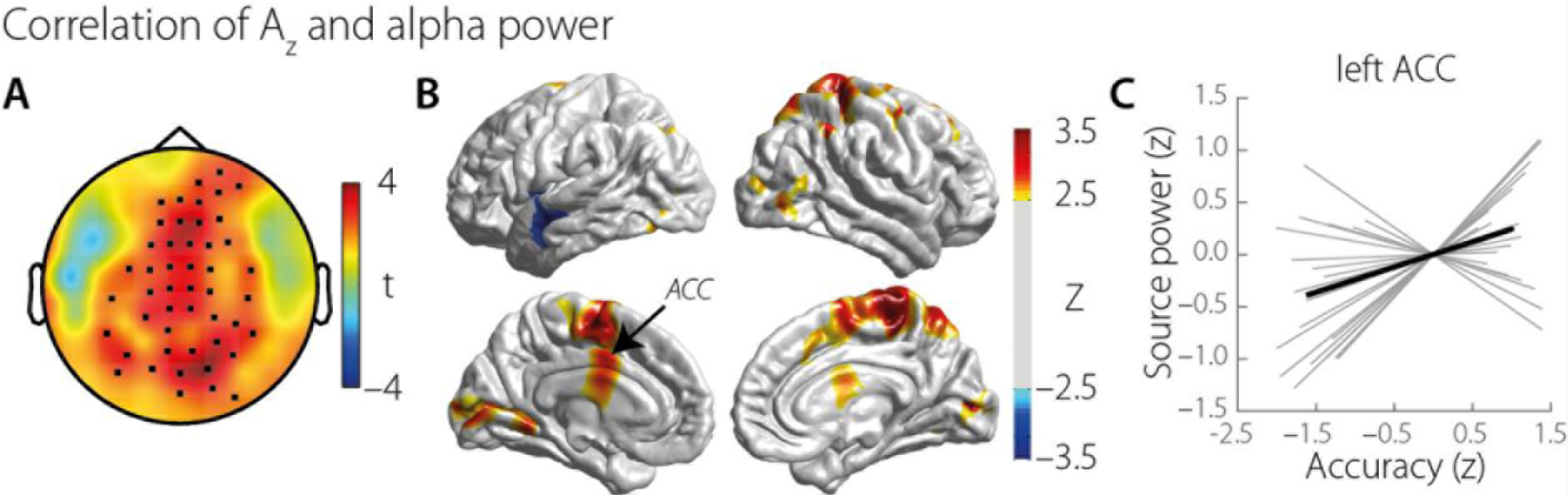
Correlation of sensitivity in memory performance (A_z_) and alpha power. **A.** Topography of the correlation of alpha power and A_z_ (t-values). Black dots display channels that belong to the significant positive cluster. **B.** Alpha power emerging from highlighted brain areas correlates with A_z_. Positive z-values indicate a positive correlation of A_z_ and alpha power. **C.** The gray lines show the single subject correlation of alpha power in left ACC and A_z_. The black line indicates slope of the correlation.

#### Functional connectivity with left STG

Source projections of alpha power revealed a pattern of brain regions susceptible to memory decay. Most prominent effects originated from left STG and bilateral visual cortices. In order to attain a better understanding of the functional role of alpha power and its different origins, we computed functional connectivity in the alpha range. Due to the strong alpha power effect in left STG (see Figure 3A) as well as its crucial role in auditory sensory memory (Sabri et al., 2004), left STG was used as a seed in a whole brain connectivity analysis. The aim of this analysis was to find brain regions that that were functionally connected with left STG, and where this connectivity was modulated by memory decay and temporal expectation.

Connectivity analyses revealed that phase locking between left STG and left V1 ([MNI: −33, −94, −14]) increased with longer delay-phase duration, whereas connectivity with right mid-temporal gyrus (MTG; [MNI: 67, −14, −16]) decreased with longer delay-phase duration (see Figure 4A). Additional statistical analyses of connectivity patterns also revealed that memory-decay-related changes in connectivity were modulated by temporal expectation. In right anterior cingulate cortex (ACC; [MNI: 1, 3, 37]) as well as in right inferior frontal gyrus (IFG; [MNI: 60, 7, 11]) connectivity with left STG increased with delay-phase duration after fixed onset times and decreased after jittered onset times (see Figure 4B).

To attain a better understanding of the increasing functional connectivity between left STG and left V1, we related the PLVs to memory performance (i.e., A_z_). We performed a median split on the PVLs for each delay-phase condition separately. Then we sorted A_z_ values according to high and low PLVs per delay phase. Finally, we contrasted high phase-locking A_z_ with low phase-locking A_z_ with t-tests. For the delay-phase durations of 1 and 2 s, memory performance did not differ between high and low PLVs (1-s delay: t(18) = 0.29, p = 0.775, 2-s delay: t(18) = −1.7319, p = 0.10). In the 4-s delay phase condition, memory performance was significantly better after low PLVs compared to high PLVs (t(18) = 2.43, p = 0.026; see Figure 5C). We performed the same analysis on the PLVs of the connectivity between left STG and right MTG. Here, memory performance did not vary between high and low PLVs at any of the delay-phase conditions (1-s delay: t(18) = −1.65, p = 0.117, 2-s delay: t(18) = −1.91, p = 0.072, 4-s delay: t(18) = −0.73, p = 0.476). Thus, the increased connectivity between STG and V1 impedes memory performance.

**Figure 5.**
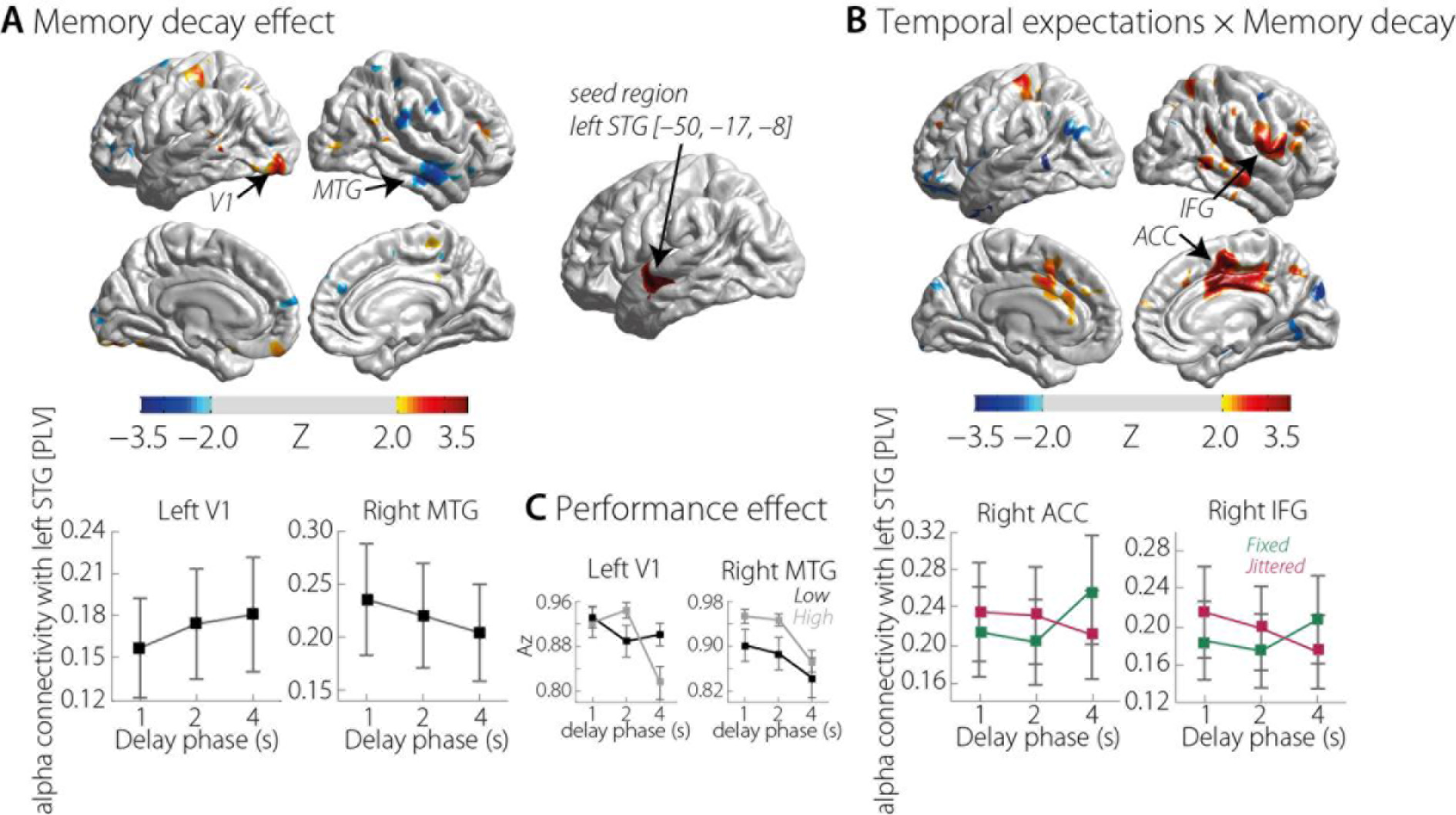
Functional connectivity. **A.** Effect of memory decay (1, 2, 4 s delay phase). Functional connectivity of left STG and highlighted brain areas is modulated by delay phase duration. Z-transformed t-values are displayed with a threshold of |z| ≥ 1.96. Positive z-values describe an increase of the phase locking value with delay-phase duration; negative z-values indicate a decrease of phase locking with delay-phase duration. Line graphs display the phase locking value between left STG and left V1 and right MTG respectively for each delay-phase duration. Error bars represent within-subject standard error. **B.** Effect of temporal expectation on memory decay. Differential impact of fixed and jittered onset times on phase locking of left STG and highlighted brain areas along different delay phases. Z-transformed t-values are displayed with a threshold of |z| ≥ 1.96. Positive z-values indicate that the slope of the correlation of phase locking and delay-phase duration is greater after fixed onset times than after jittered onset times. Negative z-values indicate that this correlation has a greater slope after jittered than after fixed onset times. Line graphs display the phase-locking value between left STG and right ACC and right IFG, respectively for each delay phase duration and each onset time condition (green line displays fixed and red line displays jittered onset times). Error bars represent within-subject standard error. The brain topography in the center illustrates the seed region (i.e., left STG) of the connectivity analysis. **C.** Effect of alpha connectivity on memory performance. Both plots show memory performance for low and high alpha connectivity between left STG and left V1 (left plot) and right MTG (right plot) for each delay-phase duration. Black lines represent performance after low connectivity, gray lines indicate performance after high connectivity. Error bars indicate standard error of the mean.

## Discussion

The current experiments assessed auditory sensory-memory decay, and showed that memory decay can be partially counteracted by temporal expectation. That is, decay is attenuated when the onset time of to-be-remembered items is fixed (and therefore highly predictable) compared to when the onset is jittered. Second, we observed a potential trading relation between alpha generated by visual and auditory regions, in that increases of alpha with delay-phase were observed in auditory cortices, while decreases were observed in visual cortices. We also observed attenuation of alpha-power modulations by temporal expectation, paralleling memory performance, in the fronto-parietal as well as the cingulo-opercular network.

### Behavioral modelling of memory decay reveals benefit from temporal expectation

In both studies, we were able to replicate the well-established finding that the longer an item is stored in sensory memory, the poorer is memory performance (e.g., Posner and Keele, 1967; Cowan et al., 1997). This can be explained by a “fading away” of the memory representation over time (Brown, 1958). Critically, in both experiments, we show that the decline of memory performance over time (decay) can be counteracted by temporal expectation. Performance was better when the onset time of the to-be-remembered sound was perfectly predictable compared to when it was jittered. Fits of an exponential decay function in Experiment 1 revealed that, for temporally predictable items, not only was decay attenuated, but it was also offset by an increase in the growth factor, which counteracts the decay factor in the exponential decay function.

Previous work suggests that prior knowledge about the time-of-occurrence of the to-be-remembered item enhances encoding precision during stimulus presentation (Rohenkohl et al., 2012), thereby allowing maintenance of the stimulus in memory for a longer period. Another (not mutually exclusive) framework, the Time-Based Resource-Sharing model (TBRS; Barrouillet et al., 2004, 2007; Barrouillet and Camos, 2012), suggests that memory traces require attentional resources to be maintained, and they decay over time as the attentional focus moves away from the representation. The higher the memory load, the fewer attentional resources are available for memory maintenance (Ma et al., 2014). We would like to suggest that temporal expectation might free attentional resources by reducing memory load (Wilsch et al., 2015b) and consequently facilitate stimulus maintenance over time.

### Differential alpha modulation in occipital and temporal cortices underlie sensory memory and mirror memory decay

Alpha power during retention was modulated parametrically by delay-phase duration. Similar to the decline in memory performance, alpha power decreased over time in bilateral primary visual cortex. Alpha-power decreases during memory delay phases have been reported to emerge from occipitoparietal brain regions (Krause, 1996; Jensen et al., 2002; Jokisch and Jensen, 2007; Tuladhar et al., 2007; Sauseng et al., 2009; Haegens et al., 2010; Bonnefond and Jensen, 2012; Wöstmann et al., 2015). Classically, occipito-parietal alpha power during auditory memory tasks is interpreted as reflecting inhibition of visual areas so that resources can be allocated to maintenance of auditory information.

In contrast, in left temporal cortex (i.e., STG encompassing primary auditory cortex), alpha power increased with longer memory-delay times. STG has been reported in previous fMRI studies to be involved during active stimulus maintenance during auditory sensory memory (Sabri et al., 2004; Grimault et al., 2009; Kumar et al., 2016). In general, recent fMRI studies indicate that activity in sensory cortices is associated with the maintenance of memory representations (for review on visual working memory, see Sreenivasan et al., 2014; for auditory cortex activity, see Linke and Cusack, 2015). Moreover, alpha power has been argued to *protect* the storage of items in memory (Roux and Uhlhaas, 2014). Corroborating this view, an auditory-memory retroactive-cueing paradigm recently demonstrated increased alpha power in a network including STG after presentation of a retro cue that allowed the participant to select an object from memory and prioritize it (Lim et al., 2015). Thus, we tentatively suggest that the observed alpha-power increase reflects the allocation of attentional resources needed to prevent the fading away of the memory representation over time, rather than inhibitory mechanisms as are classically associated with occipito-parietal alpha. Taken together, the dissociation between alpha’s behavior in visual and auditory cortices supports the presence of distributed alpha systems in the brain, employing different roles and mechanisms (Başar et al., 1997).

Finally, the alpha-band functional connectivity between left STG and left V1 increased, while connectivity between the left STG and contralateral right MTG decreased with increasing delay-phase duration. With respect to the latter finding, we suggest that the decrease in synchronization between bilateral auditory brain areas reflects in some way a lack of maintaining auditory memory representations, as different studies have shown that both left and right auditory cortices are active during auditory short-term memory (i.e., Kumar et al., 2016; Linke et al., 2015).

On the other hand, diminished V1 inhibition with longer delay-phase duration covaried with increased connection between auditory STG and visual V1, presumably allowing more interference of visual information. This interpretation is supported by our finding that lower connectivity between STG and V1 after a stimulus has been maintained for four seconds is associated with increased memory performance. Indeed, previous findings have shown that increased alpha-power connectivity can yield interference by disruptive information (in the somatosensory modality; Weisz et al., 2014) on the one hand, and that decreased connectivity can protect facial-affect recognition from disruptive visual information (Popov et al., 2013). Also, Keil et al. (2014) demonstrated that audiovisual illusory percepts were more likely to occur when connectivity between auditory and visual areas was stronger.

In sum and somewhat speculatively, the present data suggest that modulations in interregional alpha connectivity can reflect simultaneously both, gradual failure to enhance relevant processes involved in maintaining a memory representation (here, decreased STG-MTG connectivity) and failure to inhibit interfering visual activity (here, increased STG-V1 connectivity) during auditory-memory maintenance.

### The benefit from temporal expectation emerges from higher-order brain areas

One primary aim of the experiments presented here was to determine whether temporal expectation influences memory decay and the accompanying alpha power modulations. In fact, in left supramarginal gyrus (SMG) and in V1, alpha power declined faster following jittered compared to fixed onset times (similar to memory performance). For the effect in V1, we argue again that posterior alpha power functionally inhibits irrelevant information such as interfering visual input. However, the decline is less strong after fixed onset times; possibly indicating enhanced inhibition of visual information over time based on improved encoding of auditory information or sustained allocation of resources to maintenance of auditory information, as outlined above.

The SMG has previously been observed to be crucial for stimulus maintenance in auditory working memory (van Dijk et al., 2010; Obleser et al., 2012 for increased alpha power in SMG during auditory working memory; Paulesu et al., 1993). For example, Lim et al. (2015) found alpha power in SMG to be increased after a valid attention-guiding retro-cue compared to a neutral cue while maintaining a syllable in auditory working memory. In their study as well as in the present study, alpha power was increased when memory maintenance was facilitated due to an attentional cue. Furthermore, Gaab et al. (2003) investigated pitch memory with fMRI and identified left SMG to be a short-term pitch-information storage site. Notably, the BOLD signal emerging from the left SMG correlated positively with performance at the pitch-memory task, underlining the active role of left SMG for auditory sensory memory.

We also observed differential effects of fixed versus jittered onset times for alpha connectivity between left STG and the fronto-parietal network as well as the cingulo-opercular network. Connectivity increased over time in right inferior frontal gyrus; IFG and right anterior cingulate cortex; ACC. These areas are known to be relevant for top-down modulation of attention, and both regions’ BOLD activity has been shown to correlate positively with alpha power (for reviews see Dosenbach et al., 2007; Sadaghiani and Kleinschmidt, 2016). Comparable to our findings, Palva et al. (2010) demonstrated this kind of long-range communication between frontal and visual regions in visual working memory (for review, see Palva and Palva, 2011). Since these networks play an active and relevant role during the maintenance of memory representations (Postle, 2006; Jonides et al., 2008), involvement of these regions suggests that alpha-power modulations reflect active top-down modulations of STG.

Lastly, we conducted brain-wide correlations between memory performance and alpha power. We found a positive correlation emerging from anterior cingulate cortex, replicating previous findings that increased alpha power is beneficial for working memory or short-term memory performance (Haegens et al., 2010; Roux et al., 2012; Lim et al., 2015; Wilsch et al., 2015a). The anterior cingulate cortex, part of the cingulo-opercular network, is crucial for top-down control (for review, see Dosenbach et al., 2007, 2008; Petersen and Posner, 2012). Alpha power thus reflects not only an inhibitory mechanism, but appears to provide a task-beneficial ‘steering rhythm’ in and across the relevant top-down attention and sensory networks (e.g., Pinal et al., 2015).

Note that the positive correlation of performance and alpha power in V1 as well as the negative correlation emerging from left STG in this particular analysis are most likely due to the common, confounding variable of delay-phase duration itself as these regions were identified before to correlate negatively and positively with delay phase duration, respectively.

### Implications of alpha power for auditory sensory memory

Overall, the present data demonstrate how alpha power serves as a proxy for the degree of decay in sensory memory. However, the brain region in which alpha modulations are observed, as well as the direction of alpha-power changes, informs us regarding the role of alpha oscillations generated in different neural networks. Aligning our alpha-power findings with our modelling analysis of memory performance, we tentatively suggest that increased temporal alpha power after temporally expected stimuli reflects the allocation of additional resources that refresh the representation maintained in memory (Lim et al., 2015; Wilsch and Obleser, 2016). The present data show that the mechanisms by which alpha power impacts on behavioral outcomes are complex and are hardly captured by a singular mechanism, such as functional inhibition. All findings shown here, however, are compatible with a view of alpha-power as a modulatory, top-down signal (Kayser et al., 2015; Sedley et al., 2016; Wöstmann et al., 2017) that can help structure neural signaling. The present findings altogether encourage a more specific perspective on alpha power and its inhibitory role across brain areas and (trial) time. Most importantly, we were able to demonstrate that temporal expectation can alleviate memory decay, as reflected in memory performance and concomitant alpha power modulations.

## Acknowledgements

Research was supported by a Max Planck Research group grant (J.O.). The authors declare no competing financial interests.

